# Tumor localization strategies of multi-cancer early detection tests: a quantitative assessment

**DOI:** 10.1101/2024.08.07.607020

**Authors:** Christopher Tyson, Kevin H. Li, Xiting Cao, James M. O’Brien, Elliot K. Fishman, Elizabeth K. O’Donnell, Carlos Duran, Vijay Parthasarathy, Seema P. Rego, Omair A. Choudhry, Tomasz M. Beer

## Abstract

**Introduction:** Blood-based multi-cancer early detection (MCED) tests may expand the number of “screenable” cancers. Defining an optimal approach to diagnostic resolution for individuals with positive MCED test results is critical. Two prospective trials employed distinct diagnostic resolution approaches; one employed a molecular signal to predict tissue of origin (TOO) and the other used an imaging-based diagnostic strategy. Using mathematical modeling, we compared the diagnostic burden of each approach and characterized the risk of excess cancer incidence that may be attributable to radiation exposure associated with a false positive (FP) MCED test result and an imaging-based diagnostic strategy.

**Methods:** A mathematical expression for diagnostic burden was derived using MCED test positive predictive value (PPV), molecular TOO localization accuracy, and the expected number of imaging procedures associated with each diagnostic outcome. Imaging and molecular TOO strategies were compared by estimating diagnostic burden across a wide range of MCED PPVs and TOO accuracies. Organ-specific radiation dose for diagnostic imaging was extracted from the literature and used as input to National Cancer Institute RADRat tool for estimating excess lifetime cancer risk due to radiation exposure.

**Results:** For the molecular TOO diagnostic approach, an average of 2.1 procedures are required to reach diagnostic resolution for correctly-localized TPs, 4.4 procedures for incorrectly-localized TPs, and 4 procedures for FPs, vs. an average of 2.75 procedures for TPs and 2.4 for FPs with an imaging-based diagnostic strategy. Across the entire range of possible PPV and localization performance, a molecular TOO strategy resulted in a higher mean diagnostic burden: 3.6 procedures (SD 0.445) vs. 2.6 procedures (SD 0.1) for the imaging strategy. Predicted diagnostic burden was higher for molecular TOO in 95.5% of all possible PPV and TOO accuracy combinations; 79% or higher PPV would be required for a 90% accurate molecular TOO strategy to be less burdensome than imaging. The maximum rate of excess cancer incidence from radiation exposure for FP results from MCED screening between the ages of 50-84 was estimated at 64.6 per 100,000 (annual testing, 99% specificity), 48.5 per 100,000 (biennial testing, 98.5% specificity), and 64.6 per 100,000 (biennial testing, 98% specificity).

**Conclusions:** This analysis demonstrates that an imaging-based diagnostic strategy is more efficient than a molecular TOO-informed approach across 95.5% of all possible MCED PPV and TOO accuracy combinations. The use of an imaging-based approach for cancer localization can be efficient and low risk compared to a molecular-based approach.

## Introduction

Advancements in high-throughput techniques have facilitated the accurate and efficient analysis of nucleic acids and proteins in the blood, enabling sensitive detection of multiple types of cancer. As with any screening test, an abnormal multi-cancer early detection (MCED) test requires a diagnostic work-up to rule in or rule out cancer. The MCED test is unique in that it also requires the localization of the suspected tumor and the related tissue of origin (TOO).

Diagnostic evaluations are driven by clinical judgement and may include clinical examination, imaging, other diagnostic procedures, and pathologic examination of tissue. The optimal evaluation should be safe for the patient, efficient, broadly accessible, and straightforward to implement. Efficiency translates into fewer unnecessary diagnostic procedures, less cost, more rapid diagnostic resolution, and potentially fewer adverse events related to the diagnostic journey. The diagnostic evaluation must be effective in localizing a tumor and facilitating a definitive pathologic diagnosis if one is present or ruling out the presence of a tumor with a high degree of confidence to reassure the patient and avoid unnecessary anxiety.

Two prospective trials have employed distinct diagnostic resolution approaches; one employed a molecular signal to predict tissue of origin (TOO) of a suspected cancer and the other used an imaging-based diagnostic strategy. The PATHFINDER study utilized a two-step approach using: 1) molecular data obtained through MCED testing only to identify people suspected of harboring cancer, and 2) using molecular markers to predict the tissue of origin (TOO) of the suspected tumor (molecular TOO).^1^ The DETECT-A study also used a two-step process: 1) a molecular MCED test to determine those people harboring cancer, and 2) two imaging modalities (administered in a single imaging session): 1) cross-sectional anatomic imaging using IV-Contrast enhanced Computed Tomography (CT) and 2) functional imaging using CT fusion positron emission tomography (PET-CT)) to confirm and identify the location (TOO) of the suspected cancer following an abnormal MCED test result.^2^

The efficiency of a molecular TOO-informed diagnostic evaluation is challenged by several factors. Most importantly, even at very high MCED specificities, the majority of positive MCED results prove to be false positives (FPs). A recent report, for example, reported a positive predictive value (PPV) of 38% with a high specificity MCED test deployed in a predominantly elevated risk population.^1^ Patients with FP MCED results are exposed to unproductive imaging procedures, often PET-CT imaging, following an unfruitful search for a suspected cancer informed by predictive molecular TOO signal, which contributes to a high diagnostic burden. Incorrect molecular TOO predictions in patients with a true positive (TP) MCED test result, which occur in up to 15% of TP cases,^1^ further reduce efficiency, producing a similar diagnostic burden to those with a FP MCED result. Finally, comprehensive imaging is often utilized as part of the molecular TOO-informed diagnostic evaluation and even more commonly as part of tumor staging post diagnosis. Thus, patients with TP results and a correct molecular TOO prediction may gain little diagnostic efficiency from molecular TOO using current methods. To examine the risk potential of molecular TOO and compare it to an imaging approach, we modeled the anticipated diagnostic journeys and expected outcomes in a hypothetical population of average risk patients who would be eligible for MCED testing.

## Methods

Two key components were utilized to quantitatively compare the localization strategies. First, we derived a mathematical expression to calculate diagnostic burden and risk potential as a function of MCED test performance (PPV and accuracy) and diagnostic outcome. Second, we synthesized the most likely scenarios for the patient’s journey through confirmatory diagnostic options and outcomes. Clinical guidelines and relevant publications were utilized to inform the most likely course of action that a clinician might take following a positive MCED test result. The final step used the derived mathematical expression in conjunction with the clinical information to calculate our endpoints of diagnostic burden and risk potential – the former, a measure of how many procedures are required to reach diagnostic resolution, and the latter a measure of associated risk. While further MCED development may alter specific elements that we describe in this paper, we anticipate the general approach to be expandable and adaptable as the MCED field evolves.

### Mathematical Expression for Diagnostic Burden

A mathematical expression was derived from the work by Jiao *et al*.^3^ to establish a diagnostic risk potential for a TOO localization strategy as a function of the PPV of the MCED test, localization accuracy of the test, relative incidence, and risk profile of each type of cancer.

We modelled risk potential to account for all diagnostic outcomes following a positive MCED test result, incorporating a variable for the diagnostic risk potential associated with each diagnostic outcome and reframing the expression using the definition of PPV. We also considered the relative frequency (incidence) of cancer (see Supplementary Methods for full derivation).

### Assessment of Risk Potential for Each Diagnostic Outcome

The risk potential for all diagnostic outcomes was established by methodically considering the likely clinical response to the MCED test result. To estimate the diagnostic risk potential of a positive MCED result, we assessed not only the immediate diagnostic response (Table 1), but also all diagnostic procedures that would provide clinical information necessary for treatment (e.g., correct localization, pathologic confirmation, and staging of the disease). We utilized established clinical guidelines and aligned our approach with recommended diagnostic procedures across 15 common cancers of interest.^4^ We also considered the type of procedure that would most likely be used for staging and pathologically confirming the cancers of interest and compared it to the diagnostic response procedures for overlap.

**Table 1.** Cancers of interest and their associated diagnostic responses.

| **CANCER TYPE** | **DIAGNOSTIC RESPONSE** |
| --- | --- |
| BREAST | ULTRASOUND BIOPSY |
| COLORECTAL | COLONOSCOPY |
| ENDOMETRIAL | HYSTEROSCOPIC BIOPSY |
| ESOPHAGEAL | EGD BIOPSY |
| GASTRIC | EGD BIOPSY |
| HEAD AND NECK | FIBER OPTIC BIOPSY |
| KIDNEY | ULTRASOUND BIOPSY |
| LIVER | CT |
| LUNG | BRONCHOSCOPIC BIOPSY |
| NON-HODGKINS LYMPHOMA | ULTRASOUND BIOPSY |
| PANCREATIC | ERCP BIOPSY |
| PROSTATE | ULTRASOUND BIOPSY |
| SKIN | BIOPSY |
| THYROID | ULTRASOUND BIOPSY |
| URINARY BLADDER | CYSTOSCOPIC BIOPSY |

When the diagnostic response procedure would likely verify staging and/or pathologic confirmation, no further procedures would be required, but in the event the diagnostic response procedure did not fully meet these criteria, additional procedures were included.

The risk profile for each diagnostic outcome incorporated several assumptions: 1) staging and pathology cannot occur prior to localization, 2) all FPs will undergo one or more localization procedures, 3) some FPs will experience pathologic testing, 4) no FPs will experience staging procedures, and 5) disease staging requires some form of imaging, so if imaging was not part of localization or pathology, an additional imaging procedure is prescribed.

Given a sequence of procedures for each diagnostic outcome across multiple cancers, we grouped procedures into risk strata of “none,” “non-invasive,” “minimally-invasive,” and “surgery”, similar to the Lennon *et al*. DETECT-A study (Table 2).^2^ False positives and incorrectly localized TPs were assigned a null (undefined) procedure with risk equal to the mode of the correctly localized TPs. The working assumption is that the immediate diagnostic response of FPs and incorrectly localized TPs will mirror the procedures most frequently used in TPs, absent any evidence of systemic bias in TOO classifier approaches. We also considered that the negative predictive value of an organ-specific procedure in the context of ruling out cancer arising from other tissue or organs is essentially zero, and from this idea we specify that in a MCED molecular TOO strategy an incorrectly localized positive result requires subsequent imaging. This assumption is supported by reports that greater than 90% of positive results that were incorrectly localized by molecular TOO were prescribed an imaging protocol by the managing clinician.^5^ To summarize our approach for developing procedure sequences for each diagnostic outcome, we represent the procedures and assumptions for diagnostic resolution in Figure 1.

**Figure 1.**
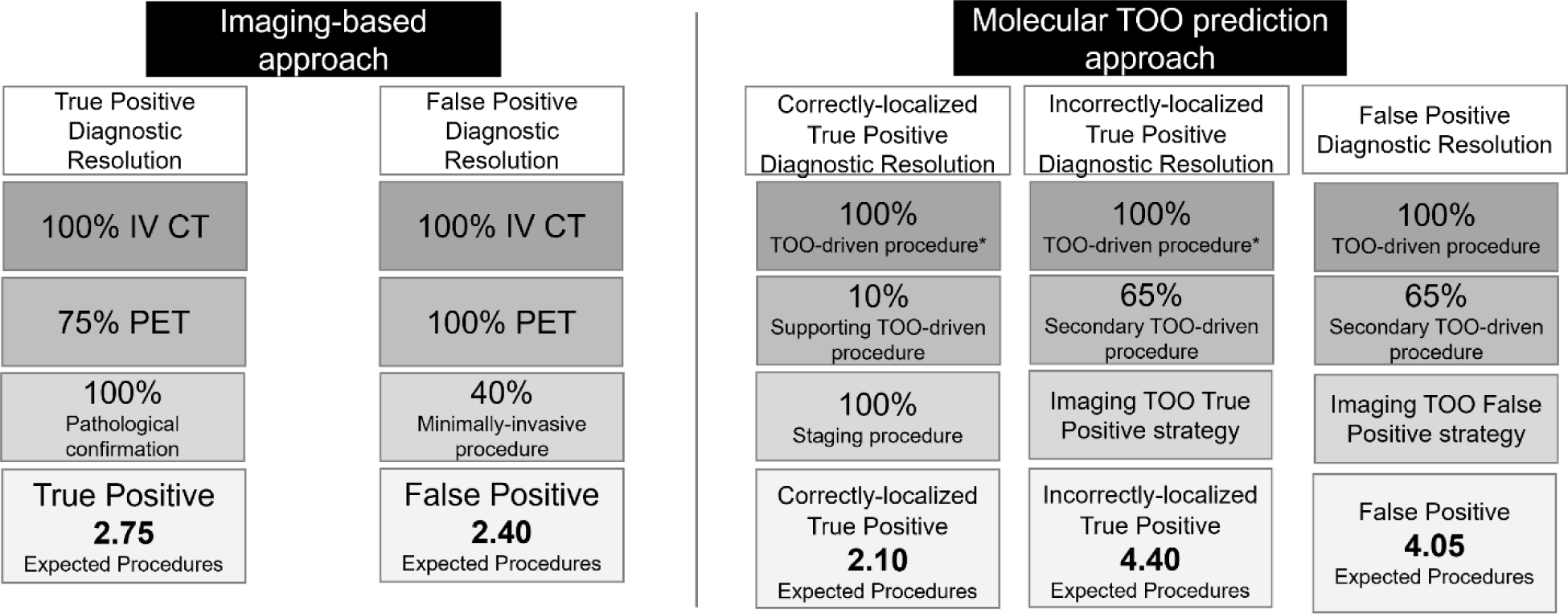
Aggregate estimates of procedures and assumptions for diagnostic resolution * Pathologic confirmation is implicitly assumed within the molecular TOO-driven procedure. CT, computed tomography; IV, intravenous. PET, positron emission tomography, TOO, tissue of origin.

**Table 2.**
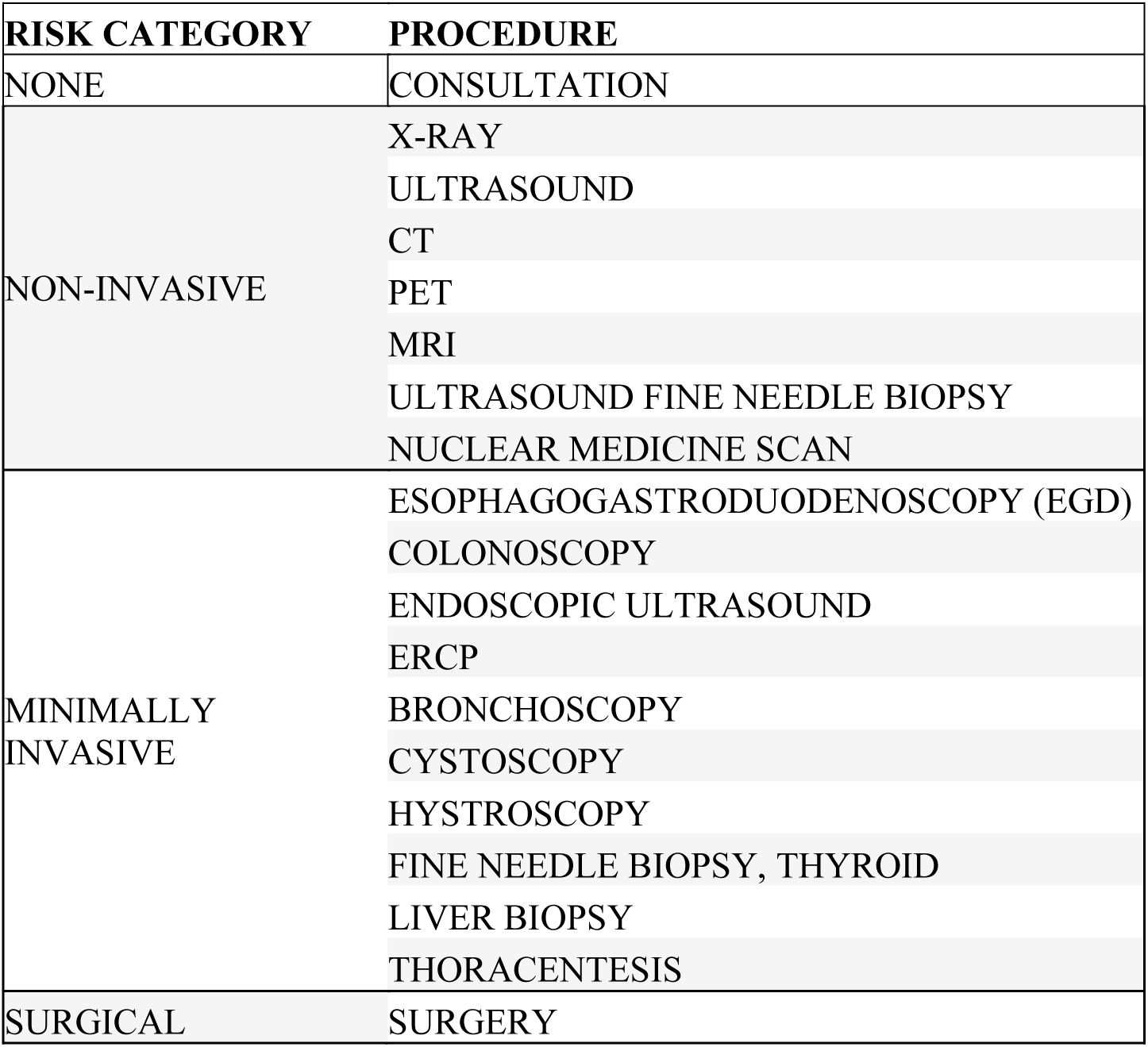
Procedures and risk stratification.

| **RISK CATEGORY** | **PROCEDURE** |
| --- | --- |
| NONE | CONSULTATION |
| NON-INVASIVE | X-RAY |
|  | ULTRASOUND |
|  | CT |
|  | PET |
|  | MRI |
|  | ULTRASOUND FINE NEEDLE BIOPSY |
|  | NUCLEAR MEDICINE SCAN |
| MINIMALLY INVASIVE | ESOPHAGOGASTRODUODENOSCOPY (EGD) |
|  | COLONOSCOPY |
|  | ENDOSCOPIC ULTRASOUND |
|  | ERCP |
|  | BRONCHOSCOPY |
|  | CYSTOSCOPY |
|  | HYSTROSCOPY |
|  | FINE NEEDLE BIOPSY, THYROID |
|  | LIVER BIOPSY |
|  | THORACENTESIS |
| SURGICAL | SURGERY |

After determining the sequence of procedures for each diagnostic outcome, we adapted the Levenshtein distance^6^ metric to quantify the risk potential for each diagnostic outcome. Qualitatively, this can be thought of as a ladder, in which the sequence begins at zero, and may take steps up (or down) the ladder. In practice, risk potential will take a positive value and then just as with a physical ladder, risk potential that is lower or “closer to the ground” is safer. Once a Levenshtein distance was calculated for each cancer type and diagnostic outcome, we used incidence rates for each cancer type from the NCI Surveillance, Epidemiology, and End Results (SEER)^7^ data to weight the risk potential of each cancer type for each diagnostic outcome and developed a cohort-level risk potential factor for each diagnostic outcome. Then we used the same mathematical formulation for diagnostic burden noted above, but instead of the number of procedures associated with each diagnostic outcome, the coefficient represents the risk potential of each diagnostic outcome.

### Estimating Radiation-Induced Cancer Risk

It is expected that radiological imaging will play a significant role in the diagnostic resolution for patients following a positive MCED test. This role may be direct, for MCED technologies that rely on radiological imaging for localization, or indirect, for technologies that rely on radiological imaging when tissue-specific confirmatory tests fail to resolve the positive MCED test result. To estimate the risk of radiation-induced cancer, a model was developed that combines the probability of exposure to a particular radiation dose, with the expected excess cancer risk attributed to that radiation dose. In this work, we focused specifically on radiological imaging sequences administered to patients following a FP test. For TP patients who are ultimately diagnosed with cancer, the radiation exposure from the radiological imaging sequence following the positive MCED test is expected to be minimal compared to the expected radiation exposure they will receive during management of their disease.

The probability of exposure for FP events was modeled for an MCED test using a Poisson process that is a function of specificity, screening interval, and screening ages (50 to 84 years old) with conservatively assumed 100% adherence (see Supplementary Methods for full derivation). We assumed independence of FP events because at present there is no available data that suggests the probability of a FP at any given screening event is influenced by results from prior screening events. We assumed 100% adherence which represents the maximum probability to experience FP events; lower adherence rates would decrease the probability of multiple FP events.

Having established the probability of experiencing FP events, the excess lifetime risk (ELR) from radiation exposure due to required radiological imaging follow-up was estimated using the National Cancer Institute’s RadRAT tool ( https://radiationcalculators.cancer.gov/radrat/). Organ—specific radiation dosing for diagnostic CTs with IV Contrast and Whole Body F18-FDG PET-CT (with use of low dose fusion CT without IV Contrast) covering the same anatomical regions under both imaging modalities was extracted from the literature (Supplementary Table S1 includes organ dosing assumptions). This data was used as input to the RadRAT simulation to estimate ELR of cancer incidence corresponding to an FP event at each year of screening eligibility.^8^ In the case of 35 annual screening events for screening from 50-84, using separate RadRAT simulations for males and females, a set of 70 estimates for ELR was produced. Using this framework, we then calculated the mean population ELR across all possible FP sequence permutations (equation in Supplementary Methods) and estimated overall population ELR that may be associated with unnecessary radiological imaging associated following a FP MCED result.

### Analyses

We performed several analyses to assess the diagnostic burden and radiation-induced cancer risk for both localization strategies. The base case scenario was evaluated using the procedure counts for each diagnostic outcome to calculate the mean and variance in diagnostic burden across the entire range of MCED test performance.

Then we performed numerical integration across the entire range of PPV and TOO accuracies (from 0 to 1) to determine the performance metrics in which each localization strategy has greater (or less) risk potential than the other strategy. Following that assessment, we broadened the approach by layering in risk potential variance for each diagnostic outcome and again using numerical integration determined the parameter space for which each localization strategy presented greater (or less) risk potential.

We also performed a probabilistic sensitivity analysis to assess the base case diagnostic burden results. The values for MCED PPV, molecular TOO accuracy, and the number of procedures were drawn from probability distributions informed by clinical guidelines. We explored nearly all possible combinations of inputs into the calculations to assess how the localization strategies compare across ten-thousand potential scenarios. Since MCED PPV is reported to range between 10% and 50%,^1,2^ we selected the median of 30% as our base scenario value for PPV. Based on the published TOO accuracy range between 80% and 95%,^1,9^ we selected a mean of 89.5% as our base scenario for TOO accuracy. We then executed the probabilistic sensitivity analysis for 10,000 iterations and computed summary statistics including median, inner quartile range (IQR), minimum, and maximum for all approaches.

A scenario analysis was also performed utilizing mean procedure counts from a prospective clinical study that used a molecular TOO localization approach.^1^ The reported mean procedure counts for the three diagnostic outcomes (correctly localized TP, incorrectly localized TP, and FP) were substituted into our model in place of the expected counts, and analyses were repeated as above.

For radiation exposure risk, we calculated the probability of a patient experiencing *k* FP events using a Poisson model that can be applied to radiological imaging utilized for diagnostic resolution following a molecular TOO prediction-informed diagnostic strategy as well as an imaging-based diagnostic strategy. We then used the described method for calculating the mean excess lifetime risk. These two approaches were combined to estimate the overall ELR at a given population level. These calculations were performed for three screening scenarios: 99% specificity performed annually, 98.5% specificity performed biennially, and 98% specificity performed biennially.

## Results

### Diagnostic Burden

We used the decision process outlined in Figure 1 to estimate the number of procedures associated with diagnostic resolution for molecular and imaging-based diagnostic strategies. Considering TP diagnostic resolution using an imaging strategy, an estimated 2.75 procedures are required, compared to 2.10 procedures (correctly localized) and 4.40 (incorrectly localized) procedures for molecular TOO localization. For FP diagnostic resolution, the number of estimated procedures was 2.40 for the imaging localization compared to 4.10 procedures for molecular TOO localization. The mean overall diagnostic burden of imaging localization was 28% lower than for the molecular TOO localization strategy, 2.6 procedures vs 3.6 procedures, respectively. A mean of 2.8 procedures for molecular localization was estimated for the scenario analysis using mean number of procedures from the clinical study^1^ as inputs to calculate estimated diagnostic burden.

When considering the entire range of MCED PPV and TOO accuracies under base case expected procedure counts, the imaging strategy resulted in less diagnostic burden than a molecular TOO strategy across 95.5% of all possible PPV and TOO accuracy combinations (Figure 2). Assuming 90% molecular TOO accuracy and the same PPV for both imaging and molecular TOO, a PPV of 79% is required for molecular TOO to have the same burden as imaging approach. When the mean number of diagnostic procedures reported in a molecular TOO based- clinical study is used in this analysis,^1^ imaging localization results in a lower diagnostic burden across all PPV and TOO accuracy combinations. This is an expected result given that the reported TP molecular-based TOO diagnostic burden was 2.8 (correctly localized) and 3.0 (incorrectly localized), which are higher than an imaging-based TP (2.75) diagnostic burden. Similarly, the molecular-based TOO FP diagnostic burden reported in the clinical study was 2.6 vs. 2.4 for an imaging-based diagnostic strategy.

**Figure 2.**
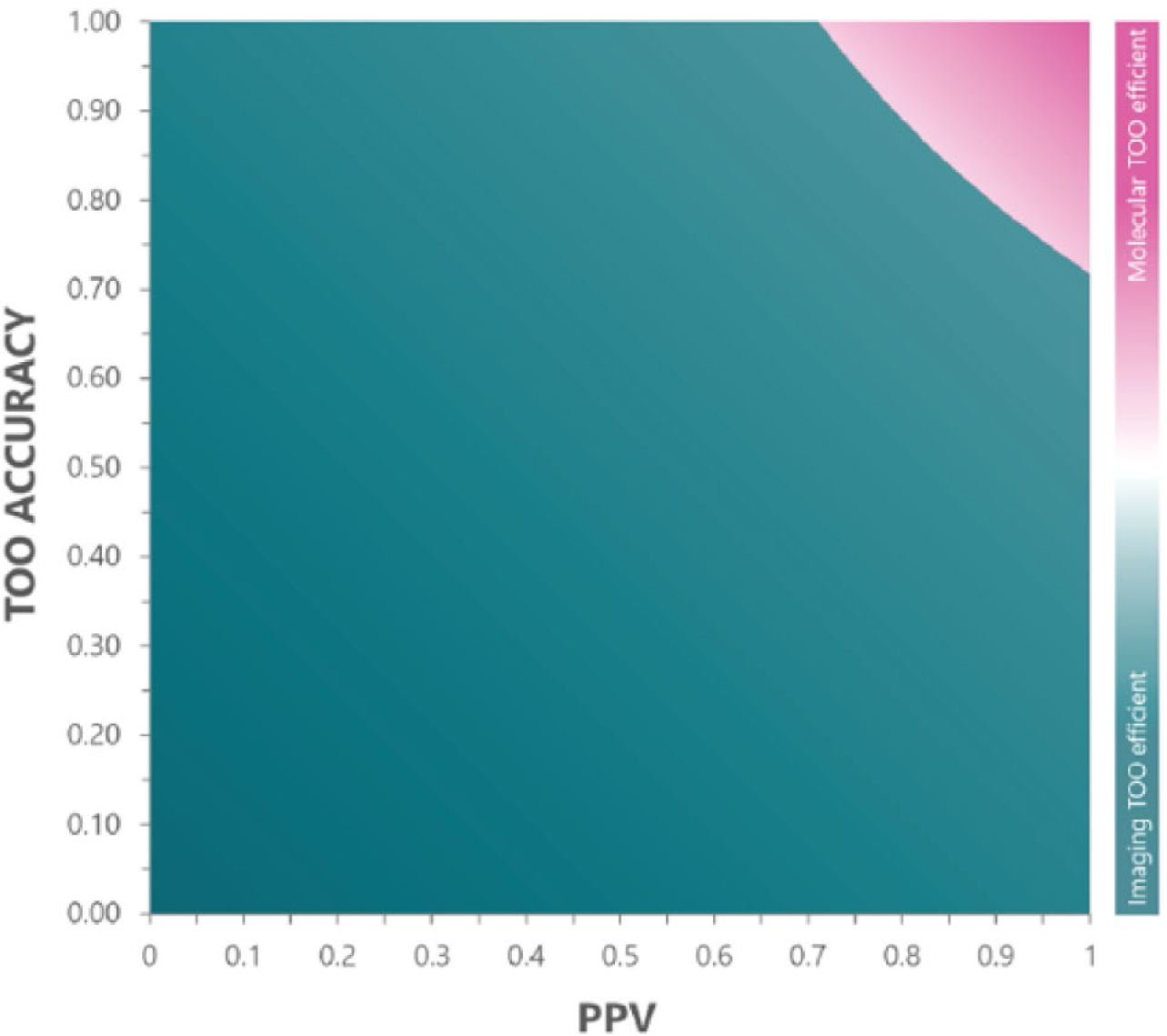
The diagnostic burden breakeven curve PPV, positive predictive value; TOO tissue of origin. The areas above and below are shaded to reflect the strategy with the lower diagnostic burden for each coordinate in the PPV and TOO accuracy plane. Areas shaded pink represent an advantage for molecular TOO, while areas shaded green represent advantage for a diagnostic imaging resolution approach.

After analyzing 10,000 probabilistic scenarios using the clinical study data, the sensitivity analysis revealed a median diagnostic burden of 2.47 procedures (IQR 0.76) for imaging localization compared to a median molecular TOO localization burden of 3.49 procedures (IQR 0.87) (Figure 3). In 97.9% of all probabilistic scenarios, a molecular TOO strategy was determined to have a higher diagnostic burden than an imaging strategy. When the molecular TOO-based clinical study procedure counts were used as inputs for a probabilistic sensitivity analysis a median burden of 2.74 procedures (IQR 1.88) was estimated.^1^ We found that 57.7% of all probabilistic scenarios were favorable for imaging TOO localization compared to molecular TOO localization using the clinical study values.

**Figure 3.**
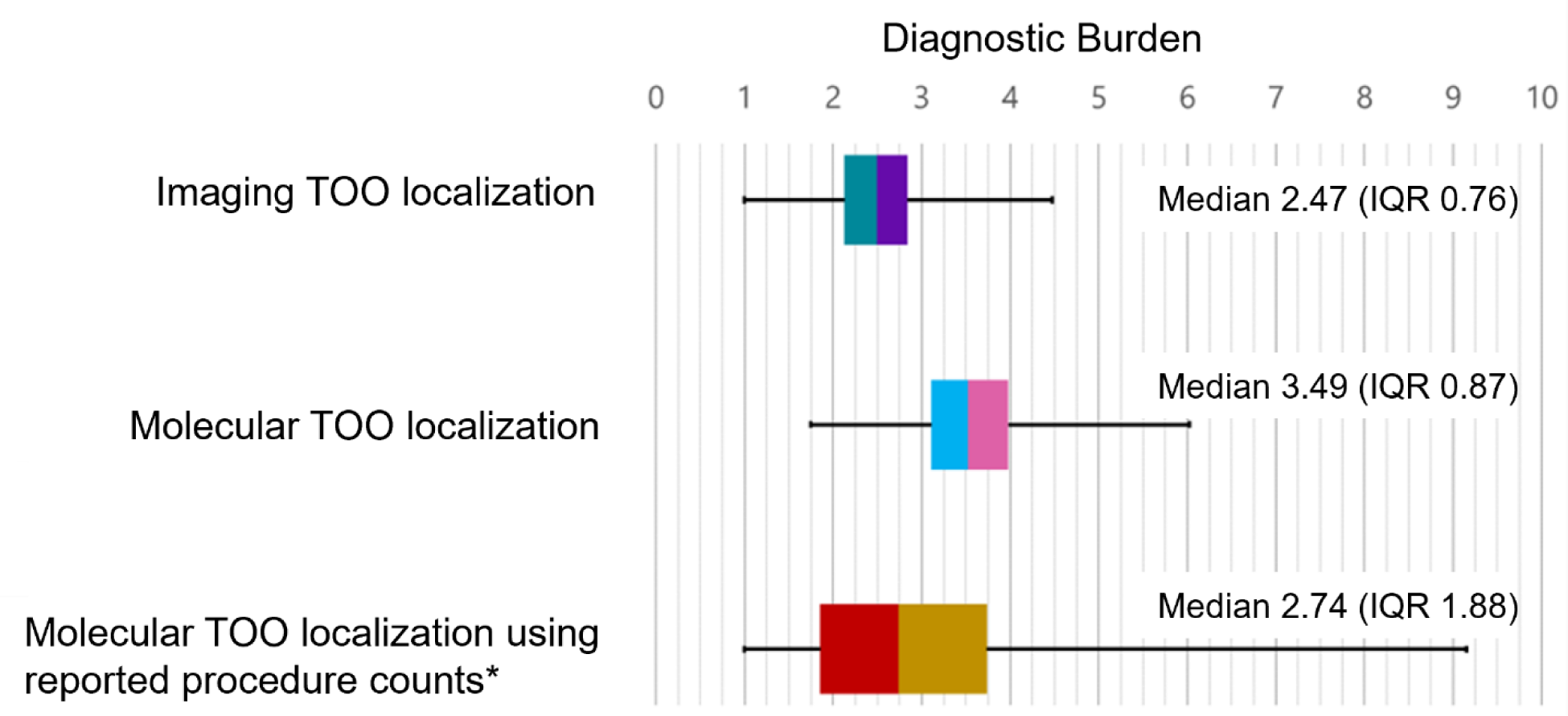
Probabilistic sensitivity analysis TOO, tissue of origin; IQR, interquartile range. *Reported procedure counts from a clinical study that utilized molecular TOO localization.^3^ The boxes represent the first quartile, median, and third quartile, while the whiskers indicate the minimum and maximum.

### Follow-up Procedure Risk Potential

The expected procedure flows for imaging- and molecular-TOO localization strategies are outlined in Figure 1. Using these procedure flows to estimate the invasive procedure risk potential of imaging localization, we derived estimates of 3.57 procedures in aggregate for TPs, and 2.96 procedures for FPs; while for molecular localization the procedure risk potential was estimated at 2.77 procedures in aggregate for correctly-localized TPs, 6.87 procedures in aggregate for incorrectly-localized TPs, and 6.26 procedures for FPs.

Using a similar approach as the diagnostic burden analysis, we then calculated the mean invasive procedure risk potential for all PPV and TOO accuracies for molecular localization to be an estimated 5.54 (StD 0.812), while for imaging localization a mean of 3.27 was estimated (StD 0.179). The risk potential breakeven curve plot (Figure 4) shows imaging localization is favorable across 98% of all PPV and TOO accuracy possibilities.

**Figure 4.**
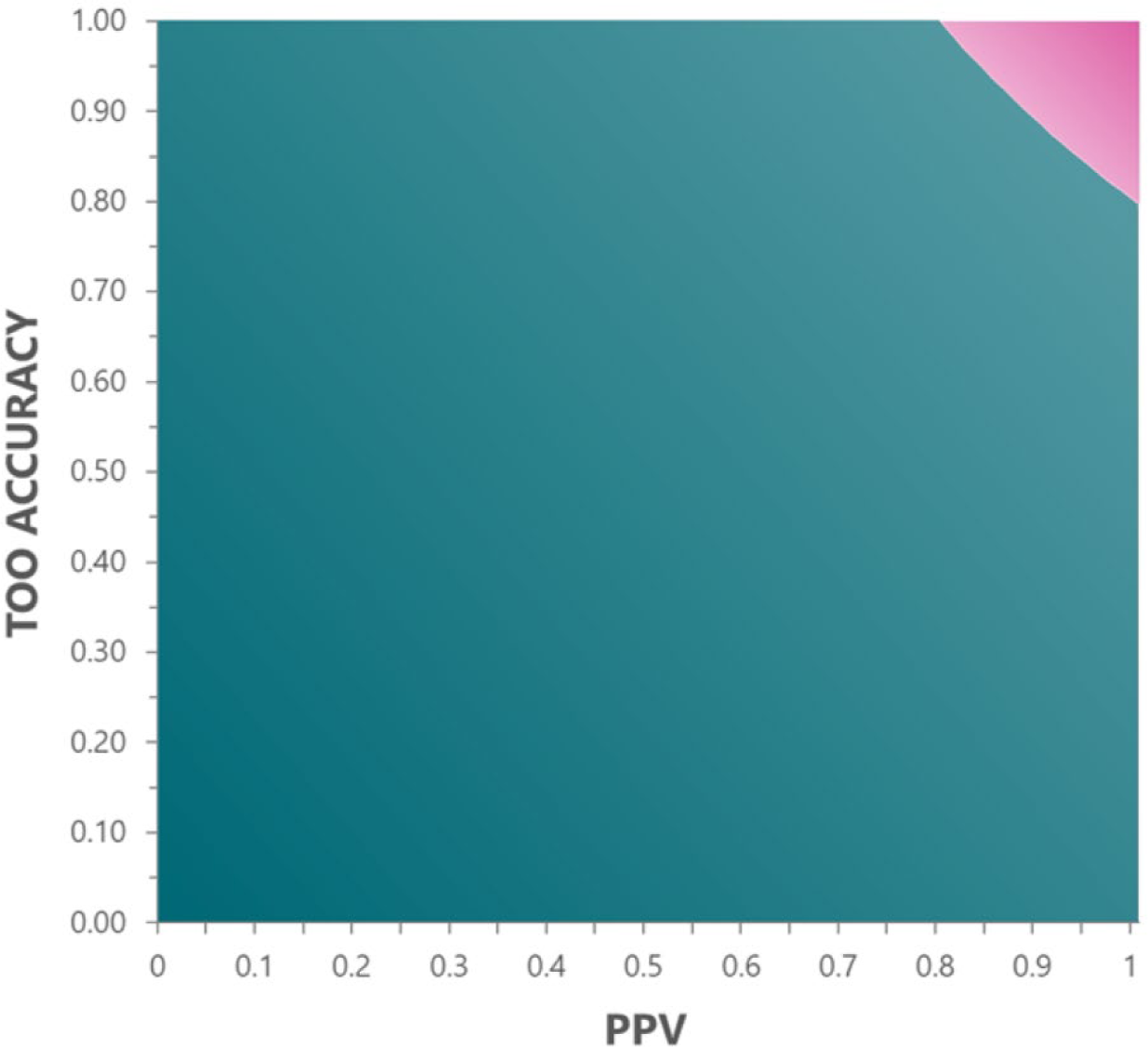
Invasive procedure risk potential PPV, positive predictive value; TOO tissue of origin. The areas are shaded to reflect the strategy with the lower invasive risk potential, with a molecular TOO strategy shaded pink, and the diagnostic imaging resolution strategy shaded green.

Subjecting the risk potential estimates to the same sensitivity approach as the diagnostic burden delivers similar results; 97.3% of probabilistic scenarios show less risk potential for imaging localization compared to molecular localization. Overall, a median risk potential of 3.15 (IQR 0.69) is calculated for imaging localization while a median of 4.14 (IQR 0.85) is calculated for molecular localization.

### Radiation Exposure Risk

Table 3 shows the probability of FPs occurring for a patient fully adherent to the specified MCED screening strategy; it is essentially the same for a molecular TOO approach and one that uses imaging for diagnostic resolution. In all scenarios, the analysis shows that at least 70% of patients will never experience any FP events and therefore never be exposed to risk from unneeded radiological imaging.

**Table 3.**
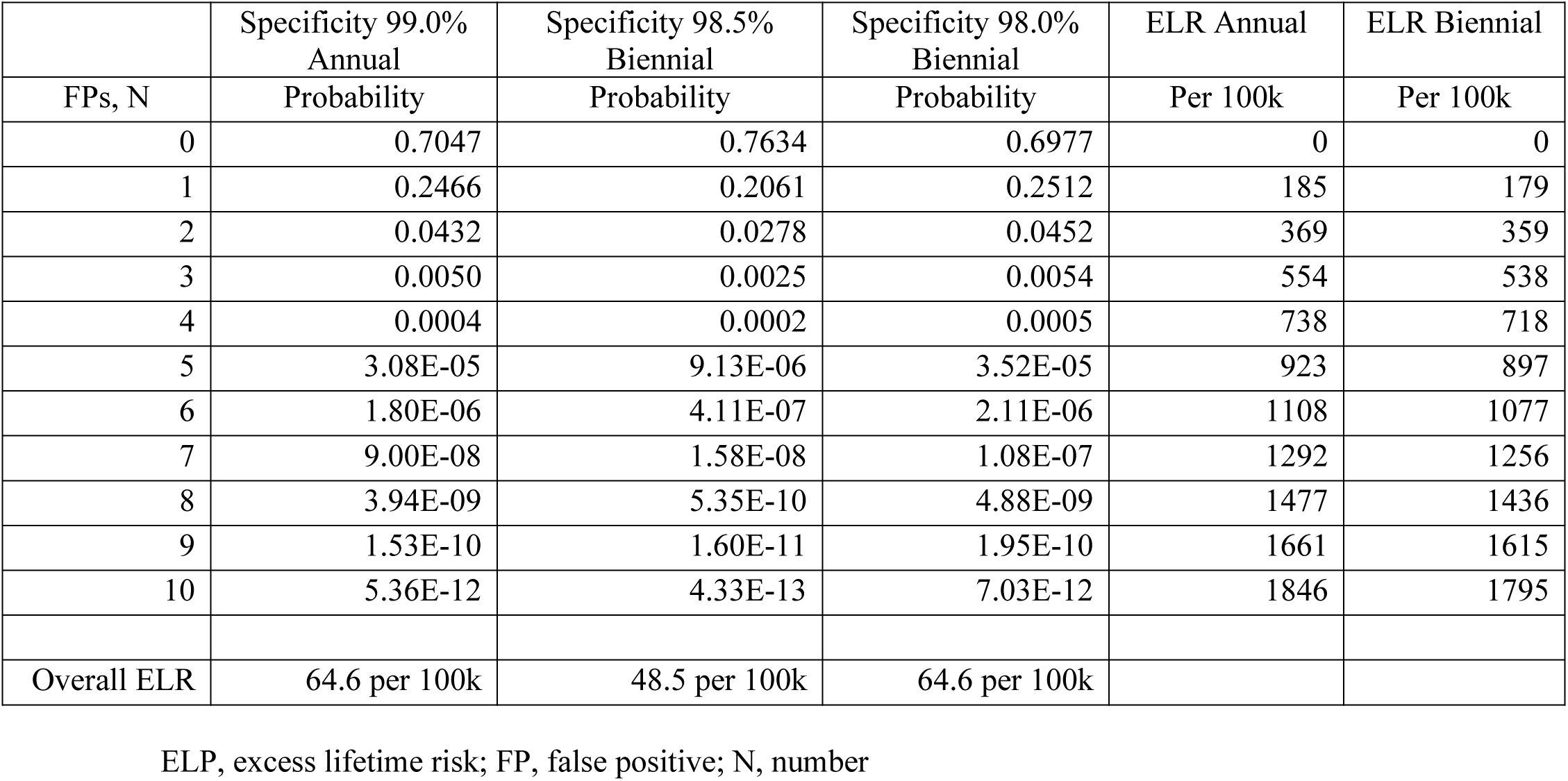
Probability of false positives occurring for a patient 100% adherent to the specified MCED screening strategy.

|  | Specificity 99.0%<br>Annual | Specificity 98.5%<br>Biennial | Specificity 98.0%<br>Biennial | ELR Annual | ELR Biennial |
| --- | --- | --- | --- | --- | --- |
| FPS, N | Probability | Probability | Probability | Per 100k | Per 100k |
| 0 | 0.7047 | 0.7634 | 0.6977 | 0 | 0 |
| 1 | 0.2466 | 0.2061 | 0.2512 | 185 | 179 |
| 2 | 0.0432 | 0.0278 | 0.0452 | 369 | 359 |
| 3 | 0.0050 | 0.0025 | 0.0054 | 554 | 538 |
| 4 | 0.0004 | 0.0002 | 0.0005 | 738 | 718 |
| 5 | 3.08E-05 | 9.13E-06 | 3.52E-05 | 923 | 897 |
| 6 | 1.80E-06 | 4.11E-07 | 2.11E-06 | 1108 | 1077 |
| 7 | 9.00E-08 | 1.58E-08 | 1.08E-07 | 1292 | 1256 |
| 8 | 3.94E-09 | 5.35E-10 | 4.88E-09 | 1477 | 1436 |
| 9 | 1.53E-10 | 1.60E-11 | 1.95E-10 | 1661 | 1615 |
| 10 | 5.36E-12 | 4.33E-13 | 7.03E-12 | 1846 | 1795 |
| Overall ELR | 64.6 per 100k | 48.5 per 100k | 64.6 per 100k |  |  |

ELP, excess lifetime risk; FP, false positive; N, number

Depending on the screening scenario, between 20% - 25% of patients may experience 1 FP event, and approximately 3% - 4% may experience two FP events. This means that only 1% of 100% adherent patients may experience more than two FP events, and the probability of experiencing more than five FP events is nearly zero.

The results from calculating the mean ELR from *k* false positive events are also presented in Table 3. For the ≥70% of patients who never experience a FP event, no additional risk of cancer is predicted. For patients experiencing one FP event, the estimated excess risk is 185 incident cancer cases per 100,000 patients fully adherent to 35 years of annual screening between the ages of 50-84 (0.185% increase), and 179 incident cancer cases per 100,000 patients fully adherent to 35 years of biennial screening between those same ages (0.179% increase). Biennial screening slightly shifts the ages of exposures higher, which leads to a reduction in the mean ELR. This phenomenon continues with the estimated mean ELR for larger number of FP events.

Combining the probability of *k* false positives with the mean ELR for *k* positives, we arrive at the overall ELR for each of the three scenarios in Table 3: 64.6 per 100,000 for 99% test specificity, annual screening, 64.6 per 100,000 for 98% test specificity, biennial screening, and 48.5 per 100,000 for 98.5% test specificity, biennial screening. The risk for both approaches is less than that of the modeled excess cancer incidence risk for annual mammography screening for women ages 40-74 of 125 per 100,000,^10^ and comparable to reported 10 years of annual low dose CT screening for lung cancer: 26 (men) to 81 (women) per 100,000.^11^ From an organ-level perspective, our analysis arrived at the following ranking from highest risk of incidence to lowest risk: bladder > lung > prostate > colon > stomach > liver > esophagus > oral cavity and pharynx > kidney > pancreas > breast > ovary > rectum > uterus > thyroid > gallbladder. While the RadRAT tool only takes radiation dose as input and does not differentiate between sources of radiation dose, the dose inputs for certain organs such as bladder and prostate may be comparatively high due to biokinetic concentration of 18F-FDG during the PET study.

## Discussion

As MCED tests advance in clinical development, the process by which a positive MCED result becomes a confirmed and actionable diagnosis is an element of the technology that requires significant attention. While many advocates for molecularly-derived TOO suggest that this method is an important aide to diagnosis following a positive MCED result, an approach that leverages radiological imaging to localize the tumor has shown promise in a clinical study as well.^2^ How these two strategies may compare and contrast in what they bring to the clinic and the benefits and risks of each require careful consideration. The work presented here is among the first to propose measures of impact to the clinic, patients, and population as a whole.

We note that a small fraction of patients may experience a more efficient diagnostic journey with the molecular TOO approach, but overall, our mathematical approach suggests that such efficiencies appear only for those patients whose cancer actually exists and is correctly localized by the MCED test. Following a MCED TOO-directed procedure, such a patient would be expected to have some radiological imaging as well and could conceivably be resolved after approximately 2 imaging procedures. But for those patients that are not correctly localized or do not have cancer, the initial MCED molecular TOO- directed procedure extends the diagnostic journey and increases procedural risk for those patients. A molecular TOO strategy therefore offers an advantage in evaluation efficiency to a surprisingly small fraction of patients which is related to the PPV of the MCED test and the corresponding TOO localization accuracy. As an example, an MCED test with 40% PPV and 85% molecular TOO accuracy will predict the correct TOO localization for a patient with a positive test 34% of the time, with the remaining 66% of patients undergoing unnecessary TOO-directed procedures. Because molecular TOO-directed procedures may often be more invasive (referenced in Table 2) than radiological imaging, those 66% of patients will likely be exposed to higher acute risk than an imaging-based diagnostic resolution.

It is observed that imaging is a component of both molecular TOO-informed and imaging-based approaches for localization, and while it does introduce the risk of radiation exposure to both approaches, our analyses show that the magnitude of that risk is minimal. A single FP event is calculated to represent on average, approximately a 0.2% increase in cancer risk over the patient’s lifetime. The probability of multiple FP events, and therefore multiple futile radiological imaging sequences, is calculated to be no higher than 5%. Taken together, the radiation risk presented by the diagnostic work-up of a positive MCED test is minimal, lower than the estimated risk of screening mammography at a population health level. While our ELR estimates are derived using data representative of an imaging-based approach, the ELR is likely to be similar to a molecular TOO approach. One or more radiological imaging procedures have been reported for some 90% of patients in clinical studies that utilized them for molecular TOO- based MCED testing.^1,9^

To date, there is no published clinical study evidence to suggest that either diagnostic approach is superior to the other. A head-to-head comparison of the molecular TOO prediction-informed and imaging-based strategies has not been investigated, but two large studies that have used these two approaches in a prospective setting have been reported.^1,2^ Direct comparisons between these two studies would be challenging, as they examined different MCED tests, enrolled different patient populations, and were activated at different centers and at somewhat different times.

A follow-up analysis of the prospective DETECT-A study suggests that an imaging-driven work-up is reliable. Of patients determined to have a FP MCED result in DETECT-A, patients with a positive MCED result, but a negative imaging evaluation, only three of ninety-eight patients were diagnosed with cancer over more than 4 years of follow-up to date. The annual incidence rate of cancer during follow-up was 1.0% (95% CI: 0.2%-2.8%).^12^ This incidence rate is comparable to the SEER annual incidence rate of 1.5% for women aged 65-74,^13^ suggesting that imaging localization is an effective strategy to resolve a positive MCED result. Furthermore, DETECT-A showed that an imaging-driven approach successfully localized all cancers that were diagnosed after a positive MCED test result across a broad spectrum of cancer sites.^2^

Considering average risk patients participating in MCED testing and current MCED test performance, there is a compelling rationale to consider an imaging first approach. Current MCED tests that have been evaluated in the general population exhibit a PPV that is impressive by comparison to single-cancer detection tests, but insufficient to enable molecular TOO to be advantageous. There are simply too many FP cases, and many would result in additional unnecessary tests when the evaluation is driven by a molecular TOO prediction. Further, an imaging-driven approach may prove simpler and easier to implement across diverse practice settings than a more complex molecular TOO informed approach. But one can conceive of a more nuanced approach to the use of molecular information to predict cancer location. It is possible that narrowly defined use cases for molecular TOO could emerge, particularly in situations where the MCED test delivers a very high PPV and the cancer of interest would have a diagnostic and staging evaluation that does not include whole-body imaging. However, our analysis suggests that significant improvements in MCED test performance are needed before a molecular TOO could be helpful, and even if achieved, the role of a molecular TOO would be quite narrowly defined. This study has some limitations. Available clinical data to inform model inputs is limited, and diagnostic pathways in the clinical setting may, in some instances, be more complex than those used in our modeling. In addition, clinical radiation exposure may vary slightly from modeled exposure reported herein. Finally, all diagnostic strategies to resolve an MCED positive result have limitations. Small cancers may be difficult to detect and localize, and incidental findings may prompt biopsies or other invasive diagnostic procedures. Other limitations will likely emerge as future studies evaluate diagnostic strategies following MCED testing. Not enough is known about these limitations to compare the performance of molecular-informed vs. imaging-based diagnostic strategy. Future studies may be able to determine if one approach or the other is more efficient with respect to localization of very small cancers or complications related to incidental findings.

## Conclusion

As MCED screening tests move closer to reality, the manner in which a positive cancer signal is diagnostically resolved represents potential risk that should be considered by all stakeholders (patients, providers, payers, regulatory bodies, and decision-makers). The two approaches currently proposed include an imaging approach that utilizes a pathway of radiological imaging, or a molecular TOO that provides a predicted organ from which the cancer signal originates using blood-based biomarkers. This work is the first attempt to quantitatively assess the differences between these approaches in both efficiency and risk.

Our quantitative evaluation demonstrates that with the performance of present generation MCED tests, the molecular TOO prediction, while conceptually attractive, cannot overcome the excess diagnostic burden it introduces in patients with false positive or incorrectly localized true positive MCED test results. Because these two groups together represent a clear majority of MCED test results, molecular TOO triggers a journey of futile, unneeded procedures including some that are invasive in nature and that likely introduce a higher risk of adverse events than radiological imaging.

While radiological imaging is relatively non-invasive, it does present a risk of radiation-induced cancers. Our assessment of the risk of radiation-induced cancers shows that at a population level this risk is similar as compared to screening mammography and for 95% of patients will represent trivial risk. In conjunction with other reports characterizing the broad utility of radiological imaging in the peri-diagnostic time even in the absence of MCED, there is growing evidence that an imaging-based diagnostic strategy for cancer localization can be efficient and low risk compared to a molecular approach of current generation MCED tests. In the future, as low-dose radiological imaging continues to improve and additional advanced imaging modalities become more available, there may be imaging-based approaches that pose even lower risk than reported in this study.

## Abbreviations

CT: computed tomography
ELR: excess lifetime risk
FP: false positive
MCED: multi-cancer early detection
PET: positron emission tomography
PPV: positive predictive value
RadRAT: National Cancer Institute (NCI) Radiation Risk Assessment Tool
SEER: Surveillance, Epidemiology, and End Results
TP: true positive
TOO: tissue of origin

## Author Contributions

*Concept and design:* Tyson, O’Brien, Beer.

*Acquisition, analysis, or interpretation of data:* Tyson, Li, O’Brien, Duran, Parthasarathy, Beer.

*Drafting of the manuscript:* All authors.

*Critical review of the manuscript for important intellectual content:* All authors.

*Administrative, technical, or material support:* Rego, Choudhry.

*Supervision:* Cao, Beer.

## Acknowledgements

Acknowledgments

Writing and editorial assistance was provided by Carolyn Hall and Feyza Sancar (Exact Sciences Corporation, Madison, WI).

## References

1. Schrag D, Beer TM, McDonnell CH, 3rd, et al. Blood-based tests for multicancer early detection (PATHFINDER): a prospective cohort study. Lancet. Oct 7 2023;402(10409):1251–1260. doi:10.1016/S0140-6736(23)01700-2

2. Lennon AM, Buchanan AH, Kinde I, et al. Feasibility of blood testing combined with PET-CT to screen for cancer and guide intervention. Science. Jul 3 2020;369(6499)doi:10.1126/science.abb9601

3. Jiao B, Gulati R, Katki HA, Castle PE, Etzioni R. A Quantitative Framework to Study Potential Benefits and Harms of Multi-Cancer Early Detection Testing. Cancer Epidemiol Biomarkers Prev. Jan 2022;31(1):38–44. doi:10.1158/1055-9965.EPI-21-0380

4. Nadauld LD, McDonnell CH, 3rd, Beer TM, et al. The PATHFINDER Study: Assessment of the Implementation of an Investigational Multi-Cancer Early Detection Test into Clinical Practice. Cancers (Basel). Jul 13 2021;13(14) doi:10.3390/cancers13143501

5. Schrag D, McDonnell CH, III, Nadauld L, et al. 903O A prospective study of a multi-cancer early detection blood test. Annals of Oncology. 2022;33:S961. doi:10.1016/j.annonc.2022.07.1029

6. Levenshtein V. Binary Codes Capable of Correcting Deletions, Insertions, and Reversals. Cybernetics and Control Theory. 1966;10(8):707–10.

7. SEER Explorer: An interactive website for SEER cancer statistics [Internet]. Surveillance Research Program, National Cancer Institute. 3/4/2024. https://seer.cancer.gov/statistics-network/explorer/

8. Quinn B, Dauer Z, Pandit-Taskar N, Schoder H, Dauer LT. Radiation dosimetry of 18F-FDG PET/CT: incorporating exam-specific parameters in dose estimates. BMC Med Imaging. Jun 18 2016;16(1):41. doi:10.1186/s12880-016-0143-y

9. Klein EA, Richards D, Cohn A, et al. Clinical validation of a targeted methylation-based multi-cancer early detection test using an independent validation set. Ann Oncol. Sep 2021;32(9):1167–1177. doi:10.1016/j.annonc.2021.05.806

10. Miglioretti DL, Lange J, van den Broek JJ, et al. Radiation-Induced Breast Cancer Incidence and Mortality From Digital Mammography Screening: A Modeling Study. Ann Intern Med. Feb 16 2016;164(4):205–14. doi:10.7326/m15-1241

11. Rampinelli C, De Marco P, Origgi D, et al. Exposure to low dose computed tomography for lung cancer screening and risk of cancer: secondary analysis of trial data and risk-benefit analysis. BMJ. 2017;356:j347. doi:10.1136/bmj.j347

12. Lennon AM, Buchanan AH, Rego SP, et al. Outcomes following a false positive multi-cancer early detection (MCED) test: Results from DETECT-A, the first large, prospective, interventional MCED study. Cancer Prev Res (Phila). May 6 2024;doi:10.1158/1940-6207.Capr-23-0451

13. Howlader N NA, Krapcho M, Miller D, Brest A, Yu M, Ruhl J, Tatalovich Z, Mariotto A, Lewis DR, Chen HS, Feuer EJ, Cronin KA SEER Cancer Statistics Review, 1975-2018. Accessed March 8, 2023, 2022. https://seer.cancer.gov/archive/csr/1975_2018/

