## Supplementary material for "Tumor localization strategies of multi-cancer early detection tests: a quantitative assessment": Supplementary Material.docx

**Supplementary Methods**

1. ***Mathematical expression for diagnostic burden***

An expression for diagnostic burden is derived as a function of overall test PPV, accuracy of the TOO call, and the number of procedures associated with each diagnostic outcome. We begin with the quantitative framework published by Jiao *et al*.^8^

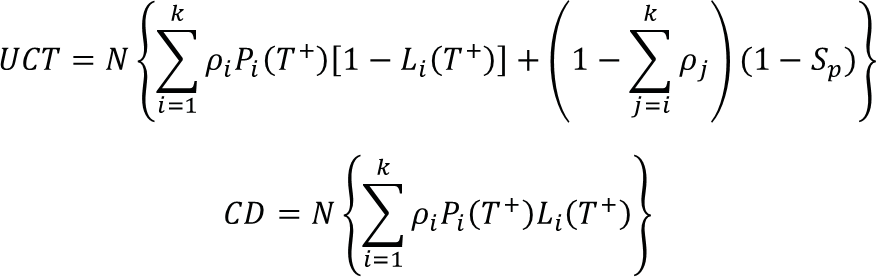

Equation 1

Equation 2

In these equations, *UCT* is the number of individuals exposed to unnecessary confirmation tests and *CD* is the number of individuals correctly identified with the disease, while *ρ_i_* is the disease prevalence for cancer *i*, *P_i_(T^+^)* represents the sensitivity of the test for cancer *i*, *L_i_(T^+^)* is the accuracy of localizing cancer *i*, and *S_p_* is the specificity of the test with respect to the set of cancers.

We arrived at an expression for diagnostic burden (Equation 3) by combining equations (1) and (2), substituting aggregate test parameters, incorporating a variable for the number of procedures associated with each diagnostic outcome, and reframing the likelihood of each diagnostic outcome using the PPV definition below:

Equation 3
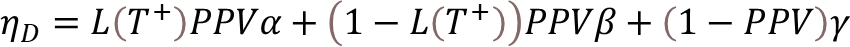

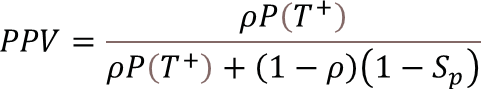
In Equation 3, *L(T^+^)* is the TOO accuracy, *α* is the number of procedures to resolve a correctly-localized TP, *ß* is the number of procedures to resolve an incorrectly-localized TP, and *γ* is the number of procedures to resolve a FP. To calculate the mean and variance for across all PPV and TOO accuracies, we employed the following characterizations:

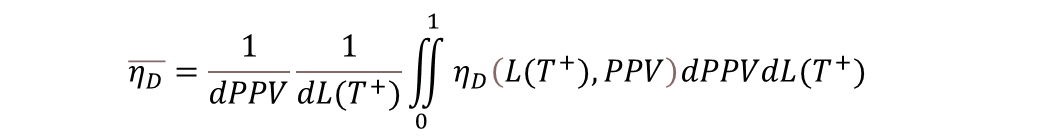

Equation 4

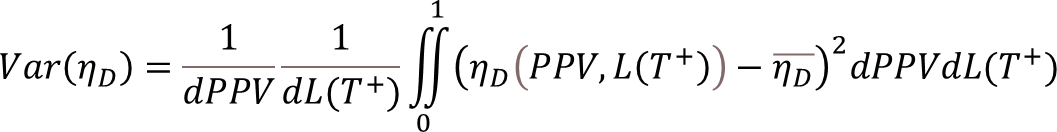

Equation 5

We also derived a breakeven equation to assess if and when one TOO localization strategy is more or less burdensome than the other. Equation 6 was used with the base case assumptions for procedure counts to calculate the break-even curve as a function of PPV.

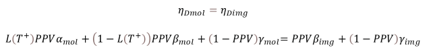

Equation 6

2. ***Estimating Radiation-Induced Cancer Risk***

Assuming that ELR for a sequence of exposures can be approximated by a linear superposition of the ELR of each element of that sequence, we derived a mathematical expression for calculating mean ELR as a function of *k* false positives over the full set of permutations. For example, for a patient that experiences FP events at ages 58, 68, and 80, we can derive the total ELR by superimposing the ELR from each single FP event. Using this framework, we can then explicitly calculate the mean population ELR across all possible FP sequence permutations.

For the final step in estimating radiation risk, we combined the Poisson process estimating the probability of *k* FPs with the mean ELR for *k* FPs to derive the overall population ELR that may be affiliated with futile radiological imaging following a FP MCED result.

First, a Poisson process was created that is a function of test specificity, testing interval, and a screening age range of 50 to 84 years old.

$$Pr\left( k;S_{p},t_{scr},a_{min},a_{max} \right)=\frac{\left[ \left( 1-S_{p} \right)\frac{a_{max}-a_{min}}{t_{scr}} \right]^{N_{FP}}e^{-\left[ \left( 1-S_{p} \right)\frac{a_{max}-a_{min}}{t_{scr}} \right]}}{k!}$$

Where *Pr(k)* is the probability of *k* false positive events, *Sp* is specificity of the blood test, *a_max_* and *a_min_* are the maximum and minimum ages of the screening paradigm, and *t_scr_* is the screening interval. Note that *a_max_, a_min_*, and *t_scr_* can be any unit of time, but must all be in the same units. In this case, the screening age range is 50 to 84 inclusive, which is 35 years.

With the probability of k false positive events determined, we must estimate the mean excess lifetime risk associated with k false positives. Provided the ELR associated with a single false positive event at a specific age, we suppose that the overall ELR induced by multiple radiation exposures due to several imaging sequences is a linear superposition of individual lifetime risks of each radiation exposure. We can then exactly and analytically solve the excess risk encompassing all permutations of false positive events across the population by noting that this is a process of set selection without repetition.

Supposing that $\left| A \right|=n$ and $a\in A$, we seek to determine how many subsets of *A* with cardinality *k* contain *a*. If we form a subset *B* of *A* such that $a\in B$ and $\left| B \right|=k$ then we must select *k­* – 1 of the other *n* – 1 elements of set *A*. This can be expressed as

$$\left( \begin{matrix} n-1 \\ k-1 \end{matrix} \right)$$

This expression represents the number of times an element a of set A appears in all permutations selecting k elements of A. For our case of screening false positives, if we wish to enumerate the number of times a false positive at 60 years of age would appear in all possible permutations of 4 false positives over 35 years of screening, we have

$$\left( \begin{matrix} 35-1 \\ 4-1 \end{matrix} \right)=\left( \begin{matrix} 34 \\ 3 \end{matrix} \right)=5984$$

Or in other words, of the 52360 ways that 4 false positives might appear when screening annually for 35 years, 5984 of those ways will include a false positive at 60 years of age. We note that in our case of selection without repetition, this value of 5984 would apply to all ages equally.

It is then clear that we can derive an expression to exactly determine the mean of all *k*-permutations of set *A* where $\left| A \right|=n$

$$\bar{A\left( n,k \right)}=\frac{\sum_{a\in A} \left( \begin{matrix} n-1 \\ k-1 \end{matrix} \right)a}{\left( \begin{matrix} n \\ k \end{matrix} \right)}$$

The numerator is the sum of all elements across all k-permutations and the denominator is the total number of k-permutations.

Supplementary Table S1. Organ-specific radiation dose (mGy) used in the ELR analyses

| Event | Organ | Male | | Female | |
| --- | --- | --- | --- | --- | --- |
|  |  | Mean | SD | Mean | SD |
| Neck CT with Contrast | Thyroid | 28.0 | 5.8 | 21.6 | 8.7 |
|  | Oral Cavity and Pharynx | 28.0 | 5.8 | 21.6 | 8.7 |
|  | Esophagus | 28.0 | 5.8 | 21.6 | 8.7 |
| Chest CT with Contrast | Lung | 20.0 | 4.2 | 15.3 | 6.2 |
|  | Breast | -- | -- | 12.0 | 4.8 |
| Abdominal and Pelvic CT with Contrast | Stomach | 18.6 | 3.9 | 14.4 | 5.8 |
|  | Colon | 17.5 | 3.7 | 13.5 | 5.5 |
|  | Rectum | 17.5 | 3.7 | 13.5 | 5.5 |
|  | Liver | 17.9 | 3.8 | 13.8 | 5.6 |
|  | Gallbladder | 18.7 | 3.9 | 14.4 | 5.8 |
|  | Pancreas | 17.2 | 3.6 | 13.2 | 5.3 |
|  | Kidney | 19.1 | 4.0 | 14.7 | 6.0 |
|  | Bladder | 18.5 | 3.9 | 14.2 | 5.7 |
|  | Uterus | -- | -- | 13.8 | 5.8 |
|  | Ovary | -- | -- | 13.8 | 5.8 |
|  | Prostate | 18.5 | 3.9 | -- | -- |
| Fusion CT without Contrast | Thyroid | 8.3 | 1.4 | 7.3 | 1.3 |
|  | Oral Cavity and Pharynx | 8.3 | 1.4 | 7.3 | 1.3 |
|  | Esophagus | 8.3 | 1.4 | 7.3 | 1.3 |
|  | Lung | 6.3 | 1.2 | 5.4 | 1.0 |
|  | Breast | -- | -- | 4.2 | 0.7 |
|  | Stomach | 5.7 | 1.0 | 4.9 | 0.9 |
|  | Colon | 5.2 | 1.0 | 4.6 | 0.8 |
|  | Rectum | 5.2 | 1.0 | 4.6 | 0.8 |
|  | Liver | 5.6 | 1.0 | 4.8 | 0.8 |
|  | Gallbladder | 5.6 | 1.0 | 4.9 | 0.9 |
|  | Pancreas | 5.2 | 1.0 | 4.5 | 0.8 |
|  | Kidney | 5.8 | 1.1 | 5.1 | 0.9 |
|  | Bladder | 5.5 | 1.0 | 4.8 | 0.8 |
|  | Uterus | -- | -- | 4.6 | 0.8 |
|  | Ovary | -- | -- | 4.6 | 0.8 |
|  | Prostate | 5.5 | 1.0 | -- | -- |
| 18F- FDG PET | Thyroid | 4.2 | 0.5 | 4.3 | 0.9 |
|  | Oral Cavity and Pharynx | 4.2 | 0.5 | 4.3 | 0.9 |
|  | Esophagus | 4.2 | 0.5 | 4.3 | 0.9 |
|  | Lung | 8.3 | 1.1 | 9.8 | 2.1 |
|  | Breast | -- | -- | 4.2 | 0.9 |
|  | Stomach | 4.7 | 0.6 | 5.4 | 1.1 |
|  | Colon | 4.9 | 0.6 | 5.8 | 1.1 |
|  | Rectum | 4.9 | 0.6 | 5.8 | 1.1 |
|  | Liver | 8.9 | 1.2 | 10.7 | 2.2 |
|  | Gallbladder | 5.4 | 0.7 | 6..0 | 1.2 |
|  | Pancreas | 5.2 | 0.6 | 6.1 | 1.2 |
|  | Kidney | 4.4 | 0.5 | 5.2 | 1.0 |
|  | Bladder | 59.1 | 3.7 | 78.9 | 9.2 |
|  | Ovary | -- | -- | 8.6 | 1.6 |
|  | Uterus | -- | -- | 8.6 | 1.6 |
|  | Prostate | 59.1 | 3.7 | -- | -- |

CT, computed tomography; ELR, excess lifetime risk; 18F- FDG PET, Flourine-18 fluorodeoxyglucose positron emission tomography; mGy, milliGray, SD, standard deviation
